# Sex differences in the association of cerebral blood flow and glucose metabolism in normative ageing

**DOI:** 10.1101/2024.11.27.625794

**Authors:** Hamish A. Deery, Chris Moran, Emma X. Liang, Caroline Gurvich, Gary F. Egan, Sharna D. Jamadar

## Abstract

It is well established that there is a local increase in cerebral blood flow and glucose metabolism in response to neuronal events. However, there is a paucity of sex disaggregated studies measuring the relationship between cerebral blood flow and glucose metabolism, despite metabolic and vascular factors being considered primary drivers of age-related cognitive decline and dementias like Alzheimer’s, which disproportionally affect women. Here we address this gap by assessing the association of cerebral blood flow and glucose metabolism in the functional networks of 79 younger and older females and males, who completed a simultaneous MR/PET scan and cognitive battery. Our results extend previously reported age-related declines in CBF and CMR_GLC_ by demonstrating that their interrelationship changes with age and sex. Older age was associated with a reduction in the correlation strength between network CBF and CMR_GLC_. CBF-CMR_GLC_ associations across people were moderated by sex, with significant negative associations in older females, a pattern not seen in older males nor younger adults. People with higher CBF-CMR_GLC_ correlations had better cognitive performance. We conclude that older adults lose synchronised vascular and metabolic dynamics in large-scale functional network, which are necessary for cognitive processes. Older females show strong, negative network CBF-CMR_GLC_ correlations, possibly reflecting a compensatory response in the face or attenuated rates of blood flow and glucose metabolism. The associations of CBF and CMR_GLC_ may serve as a biomarker for brain heath and neurological conditions.

## Introduction

The risk of cognitive decline and neurodegeneration increases in older age [1]. Although a range of risk factors have been identified, metabolic and vascular factors are considered to be primary drivers of age-related cognitive decline [2, 3]. For older women, their heightened risk of neurodegeneration is also related to hormonal changes, particularly the decline in estrogen levels that occurs during menopause [4].

Neurovascular coupling, the mechanism by which neuronal activity leads to changes in blood flow to meet the metabolic demands of the brain, was first reported in the early 20th century [5]. This discovery laid the foundation for the understanding that, during brain activation, cerebral blood flow (CBF) and cerebral metabolic rates of glucose (CMR_GLC_) and oxygen (CMRO_2_) increase [6-9]. Since this early work, neuroimaging modalities that index neurovascular coupling have been used to study changes in cerebral metabolism while people perform cognitive tasks [10]. Hybrid MR/PET scanners have been used to show that changes in glucose metabolism parallel neurovascular coupling as measured by the blood oxygenation level dependent (BOLD) response [11-14].

However, there is also heterogeneity in the association of neurovascular coupling and glucose metabolism. There is a strong spatial overlap of the BOLD signal and glucose response in the attentional networks but the well-established decrease in the BOLD signal in the default network when people perform cognitive tasks (“the task-negative response”) is not accompanied by a concomitant decease in glucose metabolism [12] (also see [15, 16]). These results highlight that the relationships that underlie neurovascular coupling can differ between brain regions and networks, and between rest and task conditions. Moreover, neurovascular coupling itself can be absent or even reversed in certain brain regions and behavioural states [10, 17], further highlighting the complex interplay between neuronal activity and the vascular, oxygen and glucose components of cerebral metabolism.

MR/PET imaging has also been used to simultaneuously index components of brain metabolism while people are in a resting state. Measuring metabolism at rest is important for understanding whether blood flow and glucose metabolism in an activated state or conditions like dementia should be attributed to the state itself or to baseline metabolic characteristics. Moderate correlations have been reported between regional CBF and CMR_GLC_ within healthy people [18-22] and between glucose and BOLD networks at rest [23], although with regional heterogeneity depending on the region’s centrality or “hubness” in the network [24, 25]. These results suggest that separate regions are perfused more or less in proportion to their metabolic demands.

It has also been suggested that the association of CMR_GLC_ and CBF across people reflects the consistency of the neurovascular coupling and that differences or changes in these associations may serve as a biomarker for brain heath [25-27]. However, reports of CBF-CMR_GLC_ associations across people have been equivocal. While positive correlations have been reported [16, 20], others have reported a lack of significant associations between CBF and CMR_GLC_ [16, 28]. We recently showed that metabolic health status can moderate the strength and direction of the association between CBF and CMR_GLC_ across people [29].

No studies to date have compared the baseline associations between resting state CBF and CMR_GLC_ in the brains of healthy younger and older females and males. This lack of research is surprising given that metabolic and vascular factors are considered to be primary drivers of age-related cognitive decline and dementias like Alzheimer’s and that females are more likely to develop Alzheimer’s than males [30]. Disruption in cerebral blood flow and glucose metabolism can lead to alterations in metabolic homeostasis and mitigate neuronal damage in dementia [26, 31, 32]. A better understanding how the association between CBF and CMR_GLC_ change in normative ageing in females and males can deepen our understanding of brain metabolism and potentially aid in early diagnosis and treatment strategies.

Here we use simultaneous MR/PET to investigate sex and age differences in association of CBF and CMR_GLC_ and cognition. We focus on functional brain networks given their importance for examining how neural information processing relates to cognition and behaviour [33]. Functional brain networks, such as the default mode, control, and attentional networks, play a fundamental role in cognition, connecting specific regions involved in complex cognitive tasks, including attention, memory, decision-making, emotional regulation and motor control [34, 35]. Disturbances in these networks in ageing or from metabolic dysfunction can lead to cognitive decline [36, 37].

Given the vascular and metabolic changes that occur in older adults and the hormonal changes that occur in females post-menopause [38, 39], we hypothesise that the association between CBF and CMR_GLC_ across individuals and across functional networks within individuals will be stronger in younger than older adults and in males than females. We also hypothesise that higher CBF-CMR_GLC_ correlations will be associated with better cognitive performance.

## Materials & Methods

### Ethical Considerations

The study protocol was reviewed and approved by the Monash University Human Research Ethics Committee. Administration of ionizing radiation was approved by the Monash Health Principal Medical Physicist, following the requirements of the Australian Radiation Protection and Nuclear Safety Agency Code of Practice (2005). For adult participants, an annual radiation exposure limit of 5 mSv applies. The effective dose in our study was 4.9 mSv. Participants provided informed consent to participate in the study.

### Participants

Ninety participants were recruited from the general community via local advertising. An initial screening interview was undertaken to confirm that participants had the capacity to provide informed consent. Exclusion criteria were a diagnosis of diabetes, neurological or psychiatric illness or dementia. Participants were also screened for claustrophobia, non-MR compatible implants, and a PET scan in the past 12 months. Women were screened for current or suspected pregnancy. Participants received a $100 voucher for participating in the study.

Eleven participants were excluded from further analyses due to excessive head motion (n=2), incomplete PET scan or blood haemolysis (n=3), incomplete ASL scans (N = 2) or consistently high or low regional ASL or CMR_GLC_ values more than 2.5 standard deviations from the mean (N = 4). The final sample included 79 individuals, 17 younger males (mean 27.5; range 20-36 years), 20 younger females (28.4; 20-42), 22 older males (76.6; 66-84) and 20 older females (75.3; 66-89) (see Table 1).

**Table 1.** Mean and standard deviation (SD) of demographics and cognitive test performance for the whole sample and the four groups based on sex and age category. p-values are for the sex, age group and sex and age group interaction effects from general linear models.

|  | Whole sample<br>(N = 79) |  | Younger males<br>(N = 17) |  | Younger females<br>(N = 20) |  | Older males<br>(N = 22) |  | Older females<br>(N = 20) |  | Sex difference |  | Age difference |  | Sex x age effect |  |
| --- | --- | --- | --- | --- | --- | --- | --- | --- | --- | --- | --- | --- | --- | --- | --- | --- |
|  | Mean | SD | Mean | SD | Mean | SD | Mean | SD | Mean | SD | p | p-FDR | p | p-FDR | p | p-FDR |
| Age (years) | 53.5 | 24.9 | 27.5 | 5.4 | 28.4 | 7.0 | 76.6 | 5.4 | 75.3 | 6.8 | NA | NA | NA | NA | NA | NA |
| Years of Education | 16.9 | 3.2 | 17.9 | 2.7 | 17.9 | 2.7 | 16.9 | 2.8 | 15.1 | 3.8 | 0.197 | 0.364 | 0.009 | 0.018 | 0.538 | 0.948 |
| Blood glucose (mmol/L) | 5.0 | 0.5 | 4.9 | 0.3 | 4.7 | 0.4 | 5.3 | 0.5 | 5.0 | 0.6 | 0.032 | 0.095 | 0.002 | 0.005 | 0.584 | 0.948 |
| Insulin (mU/L) | 4.4 | 2.9 | 6.3 | 4.2 | 4.1 | 1.5 | 4.7 | 3.0 | 3.2 | 2.3 | 0.005 | 0.023 | 0.065 | 0.091 | 0.667 | 0.948 |
| HOMA-IR | 1.0 | 0.7 | 1.4 | 1.0 | 0.9 | 0.3 | 1.2 | 0.8 | 0.8 | 0.6 | 0.005 | 0.023 | 0.119 | 0.139 | 0.711 | 0.948 |
| HOMA-IR2 | 0.6 | 0.4 | 0.8 | 0.5 | 0.5 | 0.2 | 0.6 | 0.4 | 0.4 | 0.3 | 0.005 | 0.023 | 0.345 | 0.345 | 0.924 | 0.948 |
| Systolic BP (mmHg) | 135.5 | 26.3 | 123.5 | 17.1 | 116.6 | 18.0 | 152.3 | 19.7 | 146.3 | 29.7 | 0.195 | 0.364 | 0.000 | 0.000 | 0.766 | 0.948 |
| Diastolic BP (mmHg) | 81.5 | 12.9 | 78.2 | 12.9 | 79.7 | 14.2 | 83.9 | 12.2 | 83.6 | 12.1 | 0.836 | 0.836 | 0.102 | 0.130 | 0.836 | 0.948 |
| Heart Rate (BPM) | 78.1 | 15.8 | 80.4 | 14.7 | 84.1 | 19.8 | 71.9 | 12.7 | 77.0 | 13.4 | 0.208 | 0.364 | 0.028 | 0.045 | 0.348 | 0.948 |
| Body Mass Index (BMI) | 25.0 | 4.1 | 24.3 | 3.0 | 24.4 | 5.8 | 26.3 | 2.5 | 24.7 | 4.3 | 0.415 | 0.646 | 0.237 | 0.255 | 0.482 | 0.948 |
| Cortical Thickness (mm) | 2.4 | 0.1 | 2.5 | 0.1 | 2.5 | 0.1 | 2.3 | 0.1 | 2.3 | 0.1 | 0.691 | 0.744 | 0.000 | 0.000 | 0.219 | 0.948 |
| HVLT: Delayed Recall | 8.3 | 2.6 | 9.2 | 3.0 | 9.3 | 2.0 | 6.5 | 2.3 | 8.7 | 2.0 | 0.034 | 0.095 | 0.002 | 0.005 | 0.057 | 0.399 |
| Cat Switch: Mean RT (sec) | 1.8 | 0.6 | 1.3 | 0.2 | 1.5 | 0.5 | 2.3 | 0.7 | 1.9 | 0.5 | 0.642 | 0.744 | 0.000 | 0.000 | 0.014 | 0.196 |
| Digit Sub: Correct Count | 44.4 | 22.4 | 65.3 | 9.0 | 62.7 | 14.2 | 28.5 | 11.4 | 26.7 | 16.9 | 0.473 | 0.662 | 0.000 | 0.000 | 0.888 | 0.948 |
| Stop Signal: RT (msec) | 0.56 | 0.12 | 0.53 | 0.16 | 0.52 | 0.99 | 0.60 | 0.96 | 0.58 | 0.13 | 0.539 | 0.686 | 0.029 | 0.045 | 0.948 | 0.948 |

Three participants didn't answer the years of education question; baseline blood samples not available for three participants for insulin and HOMA-IR calculations.

### Demographic and Cognitive Variables

Prior to the scan, participants completed a demographic questionnaire online. The measures included age, sex assigned at birth, gender identity, education, height and weight. Female participants reported if they were pre-, peri- or post-menopausal. Participants also completed a battery of cognitive tests, comprised of domains validated in ageing research [40] (see supplement for details): delayed recall from the Hopkins Verbal Learning Test (HVLT); reaction time in a task switching test to index cognitive control; reaction time in a stop signal task to measure response inhibition; and the number of correct responses in a digit substitution task to measure visuospatial performance.

### MR/PET Data Acquisition

Participants underwent a 90-minute simultaneous MR/PET scan in a Siemens Biograph 3-Tesla molecular MR scanner. Participants were requested to consume a high-protein and low-sugar diet for the 24 hours prior to the scan, fast for six hours and to drink 2–6 glasses of water. Participants were cannulated in the vein in each forearm and a 10ml baseline blood sample was taken. At the beginning of the scan, half of the 260 MBq FDG tracer was administered via the left forearm as a bolus. The remaining 130 MBq of the FDG was infused at a rate of 36ml/hour over 50 minutes. This combined bolus and constant infusion protocol maximises the signal-to-noise ratio over the period of the scan [41].

Participants were positioned supine in the scanner bore with their head in a 32-channel radiofrequency coil. The scan sequence started with non-functional MRI scans during the first 12 minutes, including a T1 3DMPRAGE (TA = 3.49 min, TR = 1,640ms, TE = 234ms, flip angle = 8°, field of view = 256 × 256 mm^2^, voxel size = 1.0 × 1.0 × 1.0 mm^3^, 176 slices, sagittal acquisition) and T2 FLAIR (TA = 5.52 min, TR = 5,000ms, TE = 396ms, field of view = 250 × 250 mm^2^, voxel size = .5 × .5 × 1 mm^3^, 160 slices). Thirteen minutes into the scan, list-mode PET (voxel size = 2.3 × 2.3 × 5.0mm^3^) and T2* EPI BOLD-fMRI (TA = 40 minutes; TR = 1,000ms, TE = 39ms, FOV = 210 mm^2^, 2.4 × 2.4 × 2.4 mm^3^ voxels, 64 slices, ascending axial acquisition) sequences were started. A 40-minute resting-state scan was acquired while participants watched a movie of a drone flying over the Hawaii Islands. At 53 minutes, a 5-delay pseudo-continuous arterial spin labelling (pcASL) scan was initiated (TR = 4,220 ms; TE = 45.46 ms; FOV = 240 mm; slice thickness = 3 mm; voxel size 2.5 × 2.5 × 3.0 mm^3^). Post labelling delays were 0.5, 1.0, 1.5, 2.0, and 2.5s and duration of the labelling pulse was 1.51s.

Plasma radioactivity levels were measured during the scan from 5ml blood samples taken from the right forearm, every 10-minutes, for a total of nine samples. The samples were spun in a centrifuge at 2,000 rpm for 5 minutes. 1,000-μL of plasma was placed in a well counter for four minutes and the count start time, total number of counts, and counts per minute were recorded.

### Data Preparation and Preprocessing

#### Cortical Thickness

For the structural T1 images, the quality of the pial and white matter surface was inspected, corrected where required and registered to MNI152 space. Cortical thickness for the whole brain and the Schaefer 100 regions [42] was obtained from the reconstruction statistics [43].

#### Cerebral Blood Flow

For the ASL signal quantification, a calibration map M0 of proton density weighted image was acquired for each participant. Single−subject, whole−brain CBF maps were calculated from perfusion weighted images with direct subtraction of label and control volumes. Image processing included motion, distortion and partial volume correction, a macro vascular component, adaptive spatial regularisation of perfusion and T1 uncertainty. The arterial transit time was fixed at 1.3s, T1/T1b at 1.3/1.66s, and inversion efficiency at 0.85. The resulting native space CBF images were aligned to the anatomical T1 images and normalised to MNI152 space. Grey matter CBF for the Schaefer 100 regions were obtained for each participant.

#### PET Image Reconstruction and Pre-Processing

The list-mode PET data were binned into 344 3D sinogram frames of 16s intervals. Attenuation was corrected via the pseudo-CT method for hybrid MR/PET scanners [44]. Ordinary Poisson-Ordered Subset Expectation Maximization algorithm (3 iterations, 21 subsets) with point spread function correction was used to reconstruct 3D volumes from the sinogram frames. The reconstructed DICOM slices were converted to NIFTI format with size 344 × 344 × 127 (size: 1.39 × 1.39 × 2.03 mm^3^) for each volume. All 3D volumes were temporally concatenated to form a single 4D NIFTI volume.

The PET volumes were then motion corrected [45] with the mean PET image used to mask the 4D data and corrected for partial volume effects [46], including a 25% grey matter threshold. The PET images were also spatially smoothed with surface-based smoothing [47] using a Gaussian kernel with a full width at half maximum of 8mm.

#### Cerebral Metabolic Rates of Glucose

Calculations of CMR_GLC_ were undertaken using the FDG time activity curves for the Schaefer atlas regions. The FDG in the plasma samples was decay-corrected for the time between sampling and counting as the input function to Patlak models [48].

#### Network-Level Measures

In addition to regional measures, CBF and CMR_GLC_ were calculated for the eight networks of the Schaefer atlas. The regions within the networks differ in cortical volume. Therefore, each regional value was weighted by the percentage its volume represented in the total network volume.

### Statistical Analysis

All analyses were run in SPSS version 29.0.

#### Demographics and Cognition

Univariate general linear models (GLMs) were used to test for sex, age group and sex x age group interaction effects for each demographic and cognitive measure, FDR-corrected for multiple comparisons.

#### CBF-CMR_GLC_ Correlations Across People

Whole brain and network level CBF and CMR_GLC_ associations across participants were calculated in the whole sample and in the sex and age groups separately. Two series of correlations were undertaken. In the first, no covariates were included. For the second, the demographic variables were regressed on network CBF and CMR_GLC_ separately across the whole sample. The residuals were then correlated across participants in the whole sample and for the sex and age groups separately. The correlations were tested for significance from zero using one-sample *t*-tests and between groups using z-tests, FDR-corrected for multiple comparisons.

#### Across-Region CBF-CMR_GLC_ Correlations Within Individuals

For each participant, the correlation was calculated between their 100 regional CBF and CMR_GLC_ values and between their eight network CBF and CMR_GLC_ values. Whole sample and sex and age group-averaged correlations were tested for significance from zero using one-sample *t*-tests. GLMs testing for age group, sex and age group x sex interaction effects were also undertaken.

#### Association of Network CBF and CMR_GLC_ and Cognition

Multiple regression was used to test the association between network CBF-CMR_GLC_ correlations and cognitive performance. Principle Component Analysis (PCA) was used to reduce dimensionality in the cognitive data with the resulting PC used as the dependent variable in the regression. The PCA was undertaken on normalized cognition scores and with the reaction time in the category switch and stop signal tasks multiplied by -1 so that higher scores reflect better performance in all cognitive tasks. Across-network CBF-CMR_GLC_ correlations within individuals and across-participants were used as the dependent variables. For the across-participant measure, the average network CMR_GLC_ was regressed on the average CBF value and the predicted score for each participant saved for use in the regression analysis.

## Results

### Demographic Factors and Cognition

All participants reported alignment between sex assigned at birth and gender identity. Significant sex or age group differences were found for all demographic variables, except for diastolic blood pressure and BMI (Table 1). However, none of the sex and age group interaction effects were significant at p-FDR < 0.05. All females in the older group reported being post-menopausal. Males had higher insulin, HOMA-IR than females. Older adults had fewer years of education than younger adults. Older adults also had lower heart rate and cortical thickness and higher fasting blood glucose and systolic blood pressure. Given the impact of these demographic variables on brain metabolism [29], we repeated the analyses reported below controlling for cortical thickness, blood pressure, resting heart rate, insulin resistance and BMI.

Older adults had lower HVLT delayed recall, fewer correct trials in the digit substitution task and slower reaction time in the stop signal and category switch tasks.

### Whole Brain CBF and CMR_GLC_ Associations Across People

Scatterplots of the whole brain CBF and CMR_GLC_ associations across participants are shown in Figure 1. There was a positive correlation between CBF and CMR_GLC_ that approached significance in the whole sample (r = 0.20, p = 0.077; Figure 1a.i). When cortical thickness, blood pressure, resting heart rate, insulin resistance and BMI were controlled, there was a significant negative between CBF-CMR_GLC_ correlation (r = -0.26, p = 0.028; Figure 1a.ii).

**Figure 1.**
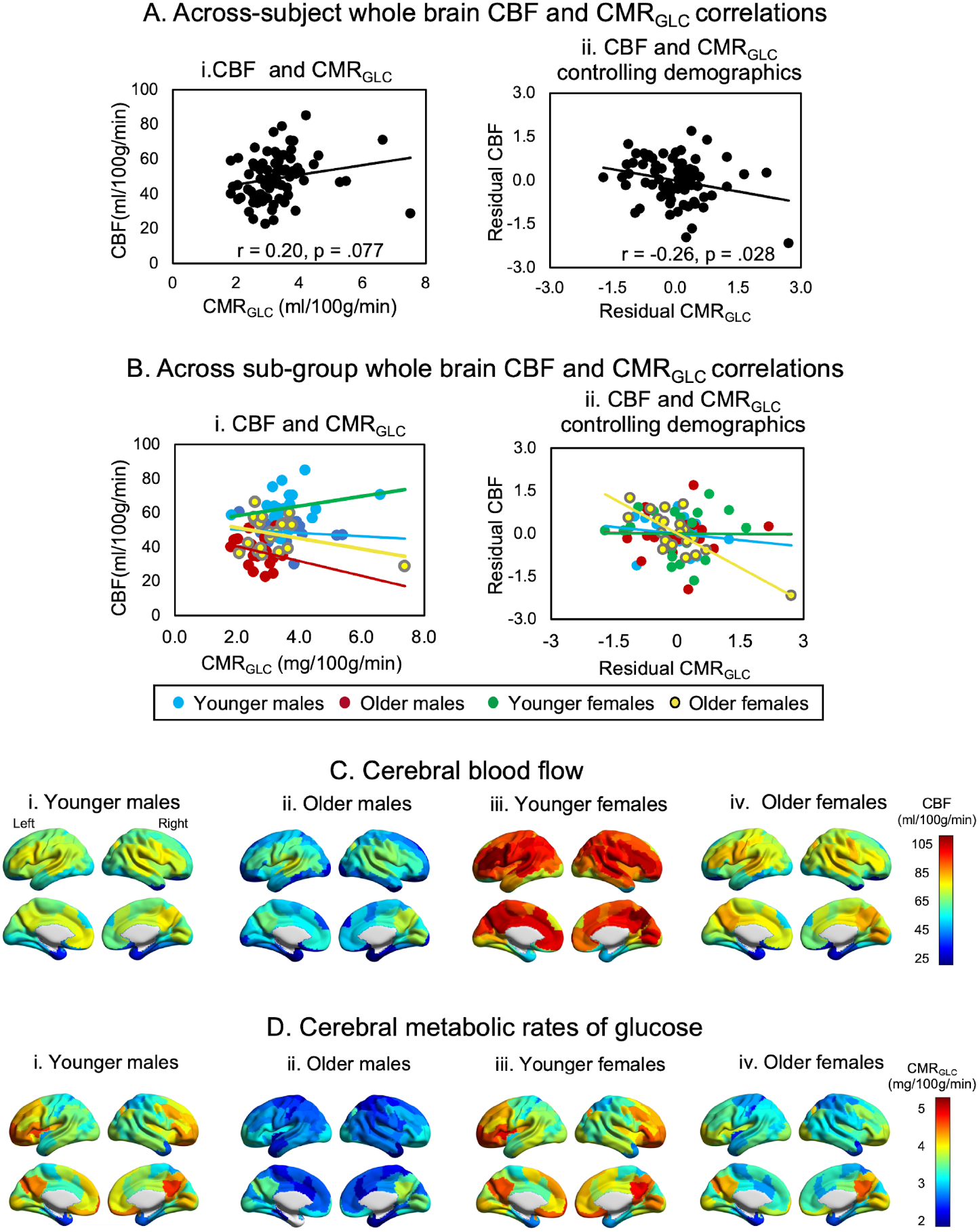
Across-subject whole brain cerebral blood flow and glucose metabolism associations. Correlation of whole brain CBF and CMR_GLC_ for the whole sample and sex and age groups with no covariates (A.i and B.i) and controlling for all demographics (A.ii and B.ii). In (A.ii and B.ii), the demographics were regressed out of CBF and CMR_GLC_ across the whole sample, before the correlations of CBF and CMR_GLC_ were calculated for each group. Regional mean cerebral blood flow (C) and glucose metabolism (D) for younger and older males and females. Significant sex and age group differences were found for CBF and CMR_GLC_ (see Supplementary Tables 1 and 2).

In the sex and age group analyses, none of the whole brain CBF-CMR_GLC_ correlations were significant (Figure 1b.i). When the other demographic variables were controlled, the negative correlation between CBF and CMR_GLC_ across older females was significant (r = -0.80, p < .001; Figure 1b.ii). The correlations among older females remained significant (r = -0.63, p = 0.004) when the participant with CMR_GLC_ above 6 ml/100g/min was excluded.

The mean regional CBF and CMR_GLC_ for the sex and age groups are shown in Figure 1C and 1D (also see Supplementary Tables 1-3). Consistent with established findings from the literature, females had higher whole brain CBF than males and higher CBF in 100 regions. Sex differences in CBF were retained in 98 regions after adjusting for the demographic variables, indicating that sex differences in cerebral blood flow are not attributable to differences in education level or the measured physiological factors (cortical thickness, HOMA-IR, blood pressure, heart rate and BMI) between the sexes.

As expected, younger adults had higher whole brain CBF than older adults and higher CBF in 97 regions. Younger adults also had higher glucose metabolism in 84 regions. None of the regional sex differences in CMR_GLC_ survived correction for multiple comparisons (Supplementary Table 2). However, the age differences in CMR_GLC_ were no longer significant after adjusting for the other physiological variables, with the effect of cortical thickness particularly strong.

### Network CBF and CMR_GLC_ Associations Across People

With the exception of the visual network, there was a positive correlation across people between network CBF and CMR_GLC_ (Figure 2a.i and Supplementary Table 5). The largest correlations were in the control (r = 0.19), limbic (0.21) and salience ventral attention (0.24) networks. The correlation for the salience ventral attention network was significant at p < 0.05 but did not survive FDR-correction. When cortical thickness, blood pressure, resting heart rate, insulin resistance and BMI were controlled, there was a negative correlation across all participants between network CBF and CMR_GLC_ (Figure 2a.ii). The largest correlations were in the default (r = -0.13), somatomotor (−0.13), visual (−0.14) and temporal parietal (−0.21) networks.

**Figure 2.**
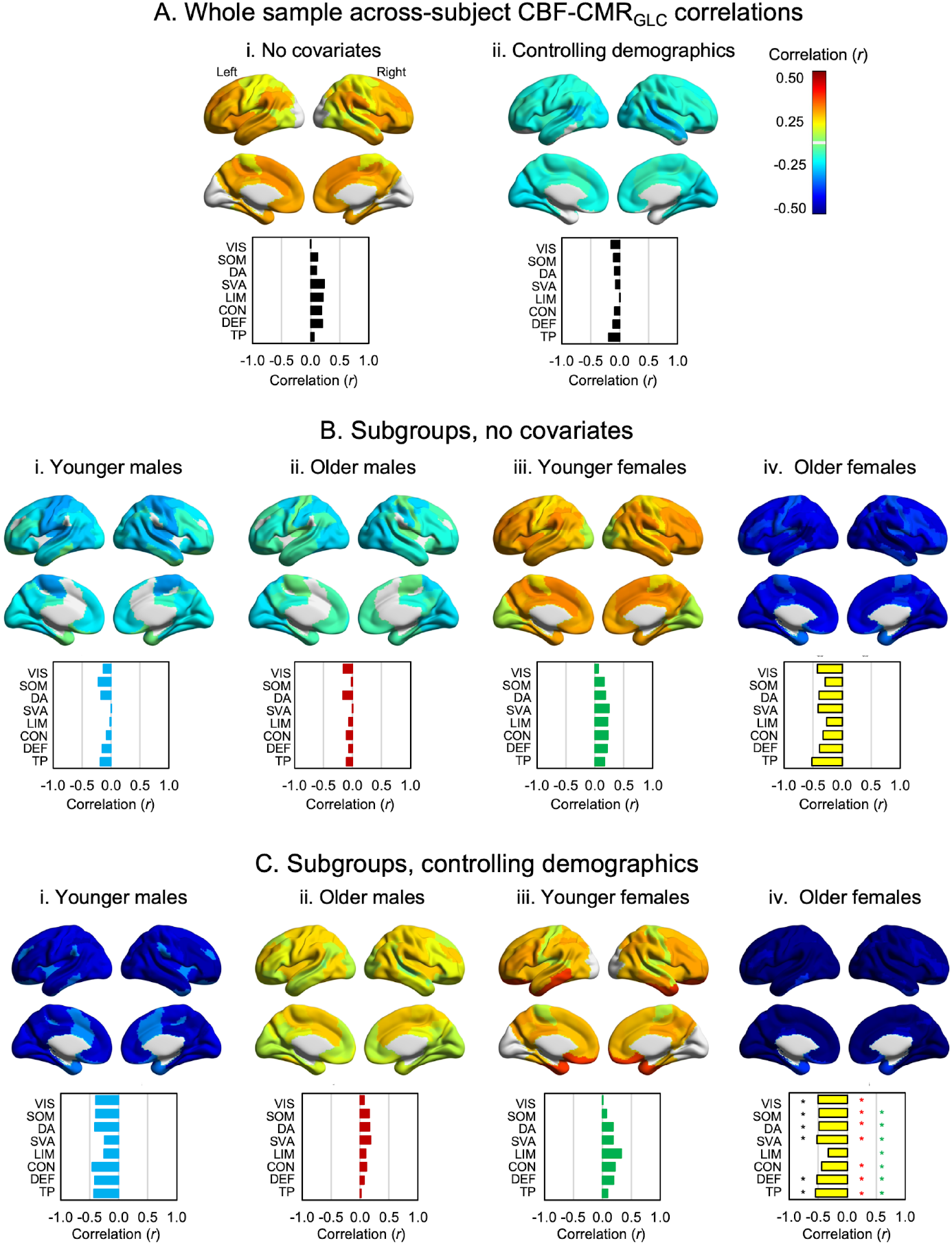
Across-subject network cerebral blood flow and glucose metabolism associations. Association of network CBF and CMR_GLC_ in the whole sample (A.i) and the sex and age subgroups (Bi to B.iv) with no covariates (and controlling for all demographics (A.ii and Ci to C.iv). For (C), black asterisk (*) to the left of the bars indicates a significant correlation different from zero at p-FDR < .05. Red asterisk (*) indicates a significant difference in correlation between older males and older females at p-FDR < .05; and green asterisk (*) a significant difference between younger females and older females at p-FDR < .05. The test of significant group differences in correlations are also summarised in Supplementary Table 5. VIS = visual; SOM = somatomotor; DA = dorsal attention; SVA = salience ventral attention; LIM = limbic; CON = control; DEF = default; TP = temporal parietal.

Across both younger and older males, small-to-moderate but non-significant negative correlations were found (Figure 2b.i and 2b.ii). Across younger females, small-to-moderate positive correlations were found between CBF and CMR_GLC_ in all networks (Figure 2b.iii). The pattern seen in younger females was inverted for older females, who demonstrated moderate-to-high negative network CBF-CMR_GLC_ correlations (Figure 2b.iv). When other demographic variables were controlled, the negative correlations across older females were significant for all networks except the limbic and control networks (Figure 2c.iv). Significant differences were also found between the correlation of older males versus older females in all networks except the limbic and control networks; and between younger females and older females in all networks at p-FDR < 0.05.

### Within Individual, Across-Network CBF and CMR_GLC_ Associations

The within-subject, across-region and across-network CBF and CMR_GLC_ correlations are shown in Figure 3 (also see Supplementary Table 6). Across the 100 regions, small-to-moderate positive correlations were found. Those correlations were significantly different to zero for the whole sample and all sex and age groups (all p < .001). Across the eight networks, the correlations were also positive and significantly different to zero for younger females (p < 0.05) and males (p < 0.01) but not older females and males. Younger adults also had higher CBF-CMR_GLC_ associations across the 100 regions and eight networks than older adults (both p < 0.05).

**Figure 3.**
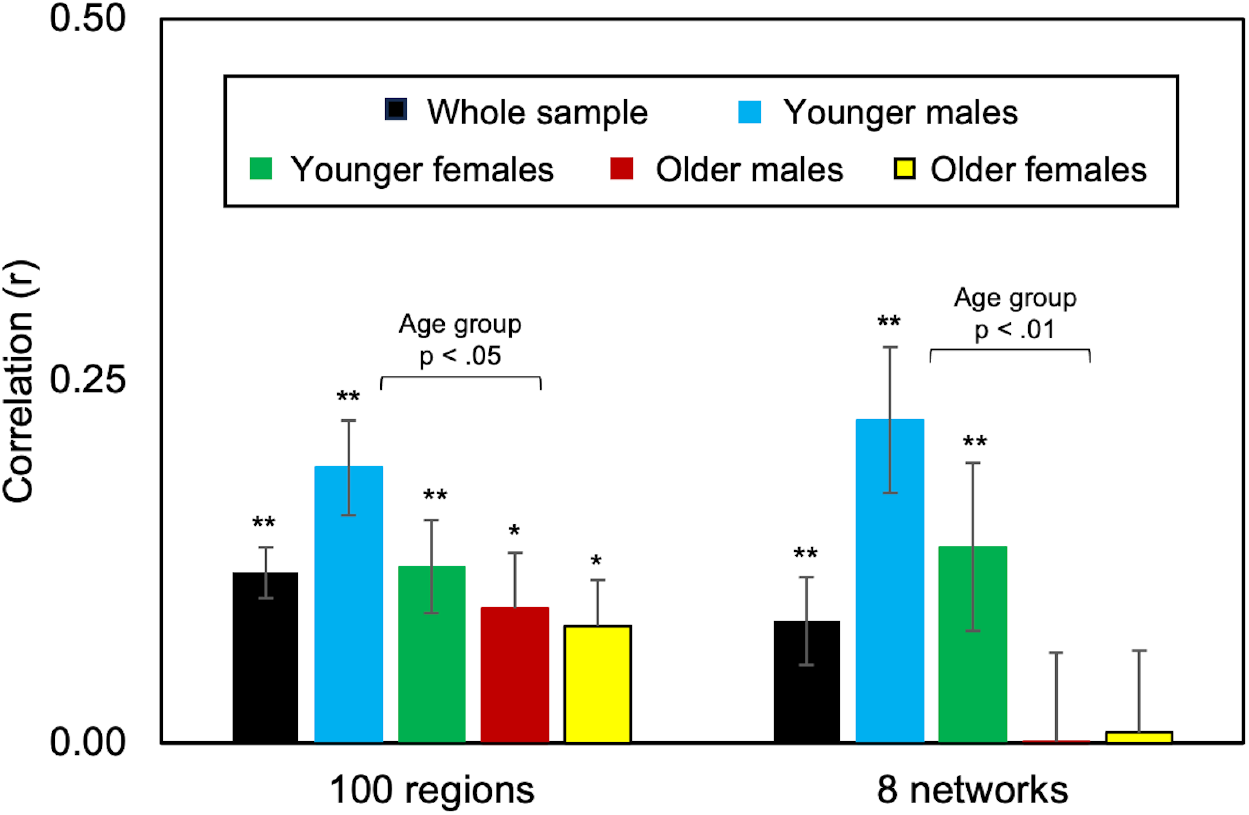
Whole sample and sex and age group mean within-subject, across-region and across-network correlations. Black asterisk indicates a correlation significantly different to zero, *p < 0.05 and **p < 0.01. The *t*-test for correlations being different to zero and GLMs of group differences are provided in Supplementary Data Table 6. The eight networks are the visual, somatomotor, dorsal attention, salience ventral attention, limbic, control, default and temporal parietal networks.

### CBF-CMR_GLC_ Associations and Cognition

One significant principal component (PC) was identified from the four cognitive measures with an eigenvalue greater than one, explaining 56.5% of the variance. The regression analysis of the association of the cognition PC with network CBF-CMR_GLC_ correlations was significant (F = 6.1, p = .004), explaining 14% of the variance. Higher across-network (beta = 0.26, p = 0.018) and across-participant (beta = 0.24, p = 0.028) network CBF-CMR_GLC_ correlations were associated with better cognitive performance (see Figure 4).

**Figure 4.**
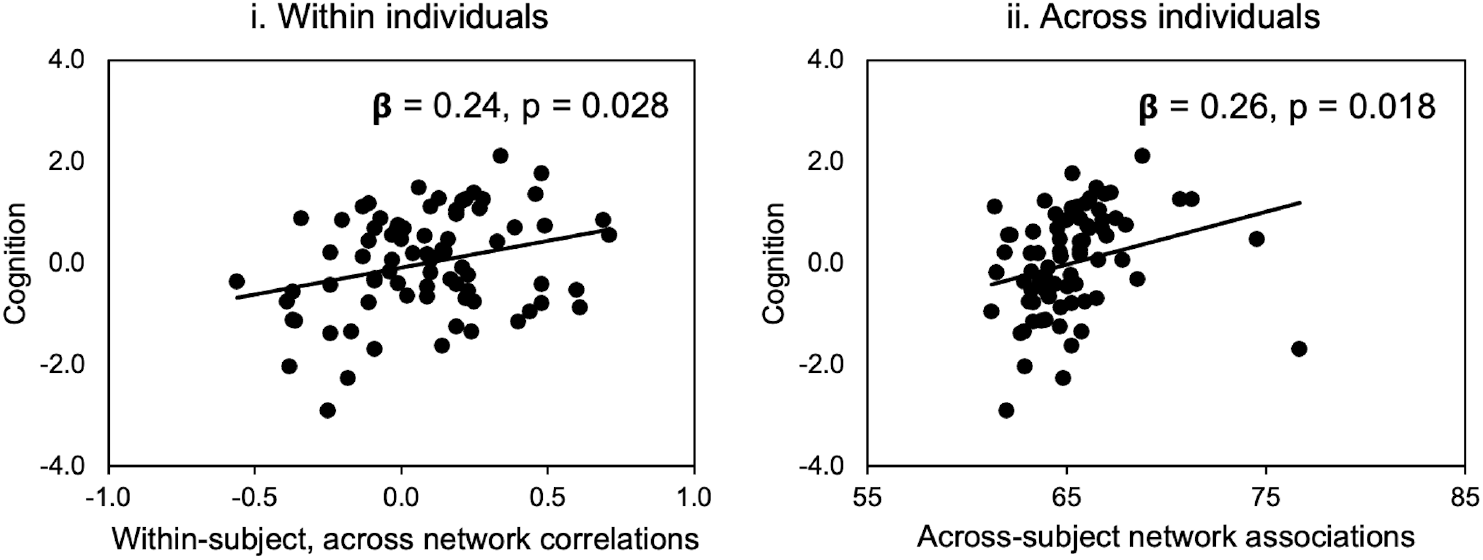
Associations of cerebral blood flow and glucose metabolism correlations with cognition. Within-subject, across networks CBF-CMR_GLC_ associations and cognition (i) and across-subject CBF-CMR_GLC_ associations and cognition (ii) from the regression analysis predicting cognition. Principle Component Analysis (PCA) of the cognitive data identified a single PC used that was used as the dependent variable in the regression analyses. Across-network CBF-CMR_GLC_ correlations within individuals and across-participants were used as the dependent variables. For the across-participant measure, the average network CMR_GLC_ was regressed on the average CBF value and the predicted score for each participant saved for use in the regression analysis.

### Anatomical Parcellation

As it is currently unknown whether there are CBF and CMR_GLC_ association differences in functional versus anatomical parcellations of the brain, we repeated the analyses using the Harvard Oxford anatomical atlas of 106 cortical and subcortical regions. Three salient differences were evident in the results of the anatomical parcellation compared to the functional parcellation (see Supplementary Information). First, the across-region CBF-CMR_GLC_ correlations were 2-3 times stronger for the anatomical than the functional parcellation (Supplementary Figure 2). Second, the CBF-CMR_GLC_ correlations across people showed similar patterns in the whole sample and sex and age groups using both parcellations, including moderate to high negative correlations among older females (Supplementary Figure 1). However, the differences between older females and the other groups were not statistically significant using the anatomical parcellation. Third, as noted above, both higher CBF-CMR_GLC_ correlations across people, and across-networks within individuals, significantly predict cognition in the functional parcellation. However, only the correlations across people significantly predicted cognition using the anatomical parcellation.

## Discussion

In this study we found a striking negative association between CBF and CMR_GLC_ across older females that was not evident across older males nor younger males and younger females. We also found that ageing is associated with a reduction in across-network correlations of CBF and CMR_GLC_ within individuals, and that people with higher CBF-CMR_GLC_ associations have better cognitive performance. Our results extend the well-documented age-related declines in cerebral blood flow [49-51] and glucose metabolism [52] by demonstrating that their interrelationship changes with age and sex, and that their relationship impacts cognition. By measuring the association of vascular and metabolic factors in functional networks, our results support the idea that brain function depends not just on absolute metabolic resource levels, but on their coordinated use across networks [34, 35]. Older adults lose synchronised vascular and metabolic dynamics in large-scale functional network, which are necessary for cognitive processes. In other words, higher perfusion and glucose metabolism in the functional networks of adults combine to fuel cognitive performance and show sex differences across people in ageing, with prominent negative associations uniquely seen in older females.

We found small-to-moderate positive correlations between CBF and CMR_GLC_ across people and across functional networks within individuals in our entire sample. The correlation coefficients were similar in strength to those reported in previous studies [21, 53, 54]. While the dynamics of resting state cerebral blood flow and glucose metabolism differ from active states, the significant correlations across networks and across people suggest that the principles of neurovascular coupling likely still apply at rest including at the spatial level of large-scale functional brain networks [6, 55]. They also support the idea that efficient neurovascular and neurometabolic function combine to support healthy brain function, providing a physiological basis for the large energy budget needed to support spontaneous brain activity at rest [56].

It has been suggested that the associations of CBR and CMR_GLC_ may serve as a biomarker for brain heath and neurological conditions [25-27]. However, to our knowledge, the utility of network CBR-CMR_GLC_ correlations for ageing and cognition has not previously been tested. Hence, our study is an important empirical step forward: it provides initial evidence that the strength of network CBF-CMR_GLC_ association – both across networks and across people – is related to age, sex and cognitive performance. If replicated and validated in larger and clinical samples, this biomarker could distinguish between normal and pathological ageing and be used for early detection before overt symptoms arise in conditions like dementia.

Although it has been known for decades that increases in neural activity drives changes in local blood flow [10], surprisingly little research has investigated sex differences in neurovascular coupling or the association of blood flow and glucose metabolism in the resting state [57]. Our results contribute to the literature by revealing that sex moderates whole brain and network CBF-CMR_GLC_ associations across people in ageing. The significant negative association between CBF and CMR_GLC_ across older females was partially consistent with our hypothesis. We hypothesised a weakening of the association due to vascular and metabolic changes that occur in ageing and the post-menopausal hormonal changes in older females [58]. We also found lower underlying rates of blood flow in older adults than younger adults. Together, these results suggest that the negative CBF-CMR_GLC_ associations in older females may reflect the brain’s drive to maintain metabolic homeostasis within a relatively tight physiological range [29]. In other words, for older females to maintain the brain’s supply of blood carrying oxygen and glucose, rates of blood flow and glucose metabolism may increase in compensation for a decrease in the other and vice versa. We also found that lower CBF-CMR_GLC_ correlations, as seen in older females, was associated with worse cognitive performance. These results suggests that the negative association of CBF and CMR_GLC_ across older females might reflect a compensatory response in an effort to support cognition in the face of age related losses of blood flow, glucose metabolism, or both.

The significant negative association between CBF and CMR_GLC_ across older females but not older males may reflect underlying biological differences, which are thought to be key contributors to different ageing trajectories and disease risks in women and men, including the risk for neurodegeneration. Changes in estrogen and testosterone contribute to the different courses of ageing in females and males (see [59] for review). Brain-derived hormones, as well as proteins and peptides that act in the brain, such as growth hormone, insulin receptor substrate 2 and insulin-like growth factor (IGF-1), also contribute to differences in brain ageing observed between the sexes [59].

To further understand the effects of age and sex on metabolism, we controlled for other physiological variables, including cortical thickness, blood pressure, resting heart rate, insulin resistance and BMI. With those variables as covariates, the number of brain regions showing age group differences in CBF was greatly reduced. Interestingly, these physiological variables did not strongly influence the relationship between sex and CBF. However, they did change the direction of correlations between CBF and CMR_GLC_ across the whole sample from positive to negative and increased the strength of the negative correlation in older females. These findings emphasise that age-related changes in cerebral metabolism are influenced by a constellation of physiological health markers, such as blood pressure, BMI and insulin resistance. While some of the apparent age-related decline in CBF may reflect poorer health, sex differences in its association with CMR_GLC_ appear more resilient to such factors, suggesting distinct biological underpinnings. Furthermore, the modulation of CBF-CMR_GLC_ coupling by health markers points to the value of integrative model of brain ageing that recognise the brain’s dependence on whole body vascular and metabolic health [60, 61].

In this study all females in the younger group were pre-menopausal (we did not control for cycle phase or use of hormonal contraceptives). All females in the older group were post-menopausal and only three (15%) were on hormone replacement therapy (HRT). Therefore, it is tempting to conclude that the differences between young and older females are hormonal in origin. While we do not have data linking sex hormones to CBF in our sample, research suggests that sex hormones play a mediating role in the control of CBF. For example, CBF and carotid artery flow increase following acute estrogen administration and long-term hormone replacement therapy in post-menopausal women [62-64]. Future research could explore these linkages by measuring sex hormones together with cerebral blood flow and metabolism.

The reason that older males do not show a similar pattern of CBF and CMR_GLC_ associations to older females is unclear. It is possible that older males compensate for a reduction in blood flow and glucose availability through different mechanisms. Reductions in CBF in older men [65] have been attributed to the modulating role of testosterone in vascular function via angiogenesis and BBB permeability [66]. It is also possible that the small, positive correlation across older males is a relative “failure” of the negative coupling seen in healthy younger males and older females. Given that older males also have lower absolute rates of CBF and CMR_GLC_ than older females, it is possible that a failure of the coupling occurs when CBF, CMR_GLC_, or both, drop below a critical threshold. Evidence for this idea comes from ischemia models in animals, in which decreases in CMR_GLC_ occur with the decrease to 38% of baseline levels, below which an increase in CMR_GLC_ is observed [67]. The possibility of a threshold for the reversal of CBF-CMR_GLC_ associations in humans, including different threshold levels in women and men, could be tested in longitudinal studies that track the time course and trajectories of changes in the rates and coupling of CBF and CMR_GLC_.

When we repeated the analyses using an anatomical atlas (see Supplementary Information) and contrasted the results to the functional parcellation, we found both similarities and differences in results. We found 2-3 times higher CBF-CMR_GLC_ across-network correlations in the anatomical parcellation, suggesting that blood flow and glucose metabolism are more tightly coupled when regions are anatomically close than when regions are distributed across space in functional networks. However, we also found that the stronger across-network correlations in the anatomical parcellation did not significantly predict cognition. We recommend that future studies continue to measure CBF-CMR_GLC_ associations based on both functional and anatomical parcellations to provide further insights into metabolic differences and to continue to test their relative utility as a biomarker of brain health and disease.

The study has a number of limitations, in particular the cross-sectional design which prevents conclusions about the causal relationship between age and cerebral blood flow and metabolism changes. As noted above, the differences found between groups may reflect unmeasured physiological, hormonal or health differences. While a key strength of this study was the simultaneous acquisition of perfusion and metabolic data, additional studies should be undertaken with larger samples. Lastly, in the absence of a middle aged cohort, we cannot infer when in the lifespan age-related changes in CBF and CMR_GLC_ and their association may occur. In particular, CBF alterations across the lifespan may follow a non-linear patterns [68]. The absence of a middle aged adult cohort in the current study precluded non-linear relationships from being tested but should be a focus of future studies.

## Conclusion

In conclusion, our results extend the previously documented attenuation of cerebral blood flow and glucose metabolism in ageing by demonstrating that their interrelationship changes with age and sex and impacts cognition. Our results support the idea that brain function depends on the coordinated deployment of metabolic substrates in functional networks. Older adults lose synchronised vascular and metabolic dynamics in large-scale functional network, which are necessary for cognitive processes. Other factors moderate this association, including sex and physiological health. Older females show a strong, negative association of blood flow and glucose metabolism across the brain, possibly reflecting a compensatory response to optimise cognition and available metabolic substrates in the face or reduced cerebral blood flow, glucose metabolism, or both. In older males, there is an absence of association. A better understanding of the interplay of sex hormones, cerebrovascular, and cerebral metabolic differences in females and males can inform the development of interventions to optimise brain function and cognition across the adult lifespan.

