## Supplementary Information for "Sex differences in the association of cerebral blood flow and glucose metabolism in normative ageing"

### 1. Cognitive Battery

The following cognitive measure were used:

**Hopkins Verbal Learning Test (HVLT).** A three-trial list learning and free recall task comprising 12 words, four words from each of three semantic categories [1]. Approximately 20–25 minutes later, a delayed recall trial and a recognition trial was completed. The delayed recall required free recall of any words remembered. The recognition trial comprised 24 words, including the 12 target words and 12 false-positives, six semantically related, and six semantically unrelated. Delayed recall (total words recalled) and a recognition discrimination index (number of correct minus number of false positives in the recognition task) were calculated.

**Digit Span.** A measure of verbal short term and working memory used in two formats: Forward and backward digit span [2]. Participants were presented with a series of digits, and are asked to repeat them in either the order presented (forward span) or in reverse order (backwards span). After two consecutive failures of the same length, the test was stopped. Scores were derived as the length of longest correct series for both forward and backward recall.

**Task Switching.** A computer-based test in which participants were given a word and had to perform one of two simple categorisation tasks, depending on the cue that appeared with the word: 1) “living” task. If the cue was a heart, participants were asked to categorise the word via a key press based on whether it represents a LIVING versus a NON-LIVING object; and 2) “size” task. If the cue was an arrow-cross, participants were asked to categorise the word via a key press based on whether it represents an object that is BIGGER or SMALLER than a basketball. The cue selection for each new trial was randomised. Half the test trials were switch trials; half non-switch trials. Half the switch and non-switch trials was congruent in the key presses for either task, half was incongruent. The measures used included the percentage of correct switch trials and mean latency of correctly responding to a switch trial [3].

**Stop Signal.** A computer-based test in which participants were presented an arrow that pointed either right or left [4]. The task was to press the left response key if the arrow pointed to the left and press the right response key if the arrow pointed to the right, unless a signal beep is played after the presentation of the arrow. In this case the response should be stopped before execution. The delay between presentation of arrow and signal beep (starting at 250ms) was adjusted up or down (by 50ms) depending on performance. The delay increased if the previous signal stop was successful (up to 1150ms) and decreased if the previous signal stop was not successful (down to 50ms). The stimulus onset asynchrony between the start of each trial (onset of fixation circles) was 2000ms. Variables were the mean reaction time in stop signal trials and stop signal reaction time. Stop signal reaction time is an estimate of inhibition ability, that is, the time required to stop the initiated go-process. The slower the stop signal reaction time, the more difficult to stop the go-process.

**Digit Symbol Substitution.** A computer-based task in which participant were presented with an 18 column x 16 row matrix [5]. The task was to translate symbols shown above the matrix (key) into digits in the matrix within a two minute period. Total count of correct responses and seconds per correct response were recorded.

### 2. Sex and Age Differences in Regional CBF and $CMR_{GLC}$

The GLMs for CBF and $CMR_{GLC}$ in the 100 brain regions are provided in Supplementary Tables 1-2. The mean regional values for each subgroup are also plotted on the brain surface in Figure 1 of the main manuscript. We found widespread sex differences across the brain in CBF with and without adjusting for the other physiological variables. We also found widespread age differences across the brain in $CMR_{GLC}$ when no covariates were included in the GLMs. However, the age differences in $CMR_{GLC}$ were no longer significant after adjusting for the other physiological variables, with the effect of cortical thickness particularly strong.

*Sex Differences:* Females had higher CBF than males in 100 regions (see Supplementary Table 3). The sex differences in CBF were largest in the lateral prefrontal cortex (effect sizes .294 to .301), intraparietal sulcus (.281 to .297), temporal cortex (.315), cingulate (.319) and precuneus (.281 to .302) in the control network; dorsal (.266 to .301) and medial (.285) prefrontal cortices in the default network; superior parietal lobule (.287 to .317), frontal eye field (.292) and post central cortex (.274) in the dorsal attention network; lateral (.316 to .336) and medial (.269 to 310) prefrontal cortices in the salience ventral attention network; and the temporal parietal cortex (.280). Sex differences in CBF were retained in 98 regions after adjusting for the demographic variables (Supplementary Table 6), indicating that sex differences in cerebral blood flow are not attributable to differences in education level or the measured physiological factors (cortical thickness, HOMA-IR, blood pressure, heart rate and BMI) between the sexes.

No effect of sex for $CMR_{GLC}$ survived correction for multiple comparisons. At the uncorrected threshold, females tended to have higher $CMR_{GLC}$ in 10 regions compared to males ($p < .05$; Supplementary Table 4).

*Age Differences:* Younger adults had higher CBF than older adults in 97 regions (p-FDR < .05; Supplementary Table 3). The age group differences in CBF were largest in the lateral prefrontal cortices (.249 to .236) and intraparietal sulcus (.247) and temporal cortex (.312) in the control network; the dorsal (.238 to 280), medial (.261 to .270) and ventral prefrontal (.265 to .282) cortices in the default network; the superior parietal lobule (.275) and frontal eye fields (.233) in the dorsal attention network; the temporal poles (.259 to .361) and orbital frontal (.260 to .319) cortices in the limbic network; the parietal operculum (.233), insula (.256) and medial posterior prefrontal (.253) cortex in the salience ventral attention network; and the temporal parietal cortex (.290). Age differences in CBF were found in 35 regions after adjusting for the demographic variables at $p < .05$, although only the age differences in the orbital frontal cortex in the limbic network survived FDR-correction (see Supplementary Table 4).

For $CMR_{GLC}$, age group differences were found in 84 regions (Supplementary Table ). Younger adults had higher $CMR_{GLC}$ in the prefrontal cortex (.166 to .168) in the control network; the dorsal prefrontal (.217 to .237), lateral prefrontal (.164) and ventral prefrontal (.172 to .206) cortices in the default network; frontal medial (.180), lateral prefrontal (.177 to .209), medial prefrontal (.211 to .274) cortices and insula (.192 to .216) in the salience ventral attention network; and somatomotor regions (.160 to .186). Age differences in $CMR_{GLC}$ were found in 39 regions after adjusting for the demographic variables at $p < .05$, although none of the regional age survived FDR correction (see Supplementary Table 7). The effect of cortical thickness on $CMR_{GLC}$ was significant in 41 regions at $p < .05$ and 26 regions after FDR-correction.

We did not find sex by age group interaction effects for regional CBF nor $CMR_{GLC}$ at p-FDR < .05 (Supplementary Tables 1 and 2).

### 3. Anatomical Parcellation

#### 3.1 Across-Subject Network CBF and $CMR_{GLC}$ Associations

Similar to the functional parcellation, there was a positive correlation across participants between CBF and $CMR_{GLC}$ (Supplementary Figure 1a.i and Supplementary Data Table 7). The largest correlations were in the frontal (r = 0.25), parietal (0.18) and motor (0.16) regions. None of the correlations were significantly different to zero with and without controlling for the demographics.

Across younger females, small-to-moderate positive correlations were found between CBF and $CMR_{GLC}$ in all areas (Supplementary Figure 2b.iii). Across both younger and older males, small-to-moderate but non-significant negative correlations were found. The pattern seen in younger females was inverted for older females, who demonstrated moderate-to-high negative correlations between network CBF and $CMR_{GLC}$. Unlike the functional parcellation, none of the differences in correlations between groups were significant at p-FDR < 0.05 with and without controlling for the demographics.

#### 3.2 Across-Network CBF and $CMR_{GLC}$ Associations

The within-subject, across-network CBF and $CMR_{GLC}$ correlations are shown in Supplementary Figure 2 (also see Supplementary Data Table 8). Across the 106 anatomical regions, small-to-moderate positive correlations were found. Those correlations were significantly different to zero for the whole sample and all subgroups (all p < 0.001). Across the eight networks, the correlations were also positive and significantly different to zero for the whole sample and all subgroups (p < 0.001). Females had lower CBF-$CMR_{GLC}$ associations across the 106 regions than males (p < 0.05).

The other salient feature of the across-space CBF and $CMR_{GLC}$ relationships is the higher correlation coefficients for the anatomical than the functional parcellation (Supplementary Figure 2).

#### 3.3 CBF-$CMR_{GLC}$ Associations and Cognition

The regression analysis of the association of the cognition PC with the anatomical CBF and $CMR_{GLC}$ associations was significant (F = 5.0, p = 0.009), explaining 12% of the variance. Higher across-subject (beta = .28, p = 0.010) CBF-$CMR_{GLC}$ correlations were associated with better cognitive performance, but the association of cognition with across-space correlations in the anatomical parcellation was not significant (beta = .21, p = 0.056).

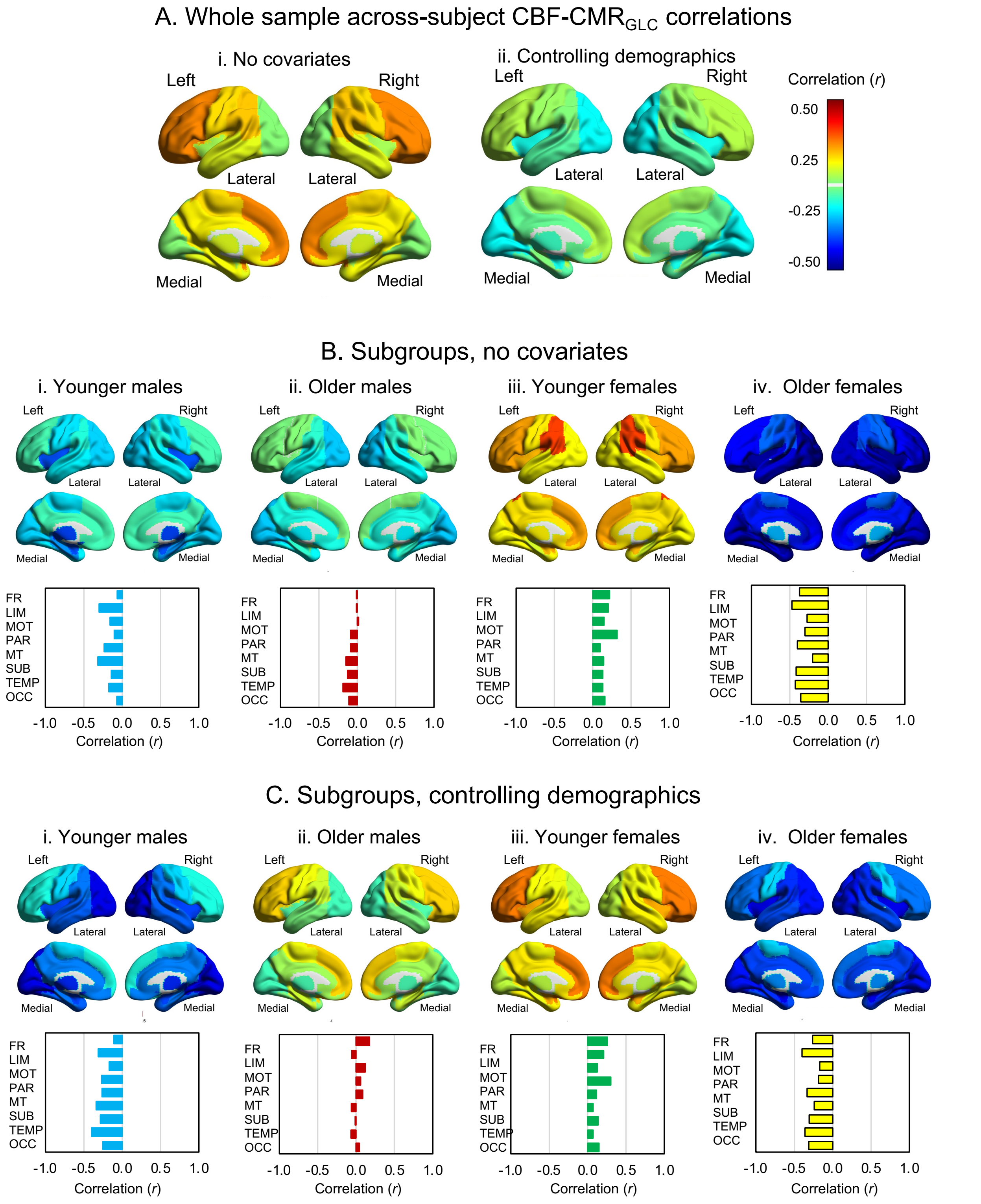

Supplementary Figure 1. Across-subject CBF and $CMR_{GLC}$ associations using Harvard Oxford atlas anatomical parcellation. Association of cerebral blood flow and $CMR_{GLC}$ in the whole sample and the sex and age subgroups with no covariates (A.i and B) and controlling for all demographics (Aii and C). None of the age group, sex or age group x sex effects were significant at p-FDR < .05.

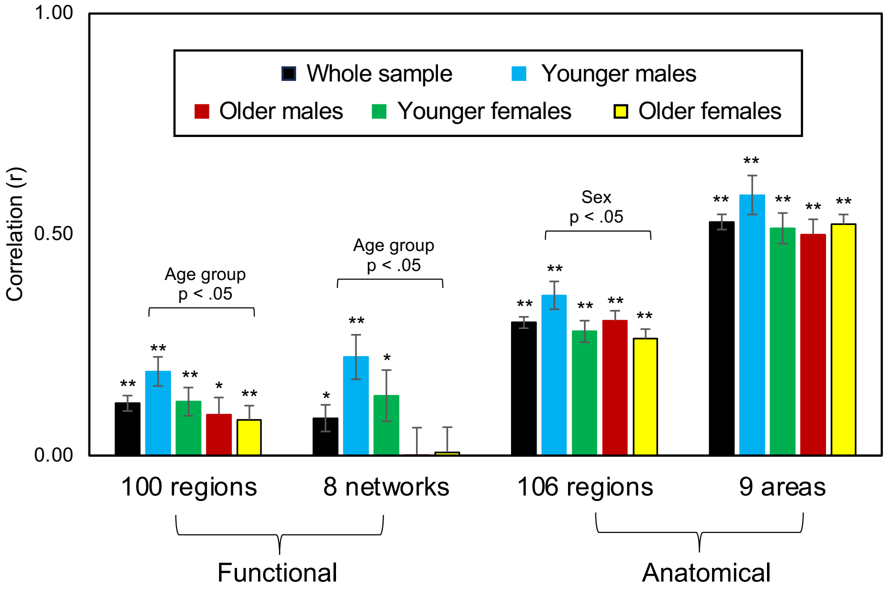

Supplementary Figure 2. Across-space CBF and $CMR_{GLC}$ associations. Whole sample and sex and age group-mean within-subject, across-region and across-network correlations for the functional parcellation reported in the main manuscript and the anatomical parcellation. Black asterisk indicates a correlation significantly different to zero, *p < 0.05 and **p < 0.01 (see Supplementary Data Table 8). Males also had higher across-space correlations than females across the 106 regions (p < .05). The eight functional networks are the visual, somatomotor, dorsal attention, salience ventral attention, limbic, control, default and temporal parietal networks; the nine anatomical areas are the frontal, limbic, motor, parietal, medial temporal, subcortical, temporal and occipital lobes/areas.
